# Neuropilin 2 signaling mediates corticostriatal transmission, spine maintenance, and goal-directed learning in mice

**DOI:** 10.1101/659342

**Authors:** Maxime Assous, Edward Martinez, Carol Eisenberg, Aleksandra Kosc, Kristie Varghese, Diego Espinoza, Shaznaan Bhimani, Fulva Shah, James M. Tepper, Michael W. Shiflett, Tracy S. Tran

## Abstract

The striatum represents the main input structure of the basal ganglia, receiving massive excitatory input from the cortex and the thalamus. The development and maintenance of cortical input to the striatum is crucial for all striatal function including many forms of sensorimotor integration, learning and action control. The molecular mechanisms regulating the development and maintenance of corticostriatal synaptic transmission are unclear. Here we show that the guidance cue, Semaphorin 3F and its receptor Neuropilin 2 (Nrp2), influence dendritic spine maintenance, corticostriatal short-term plasticity, and learning in adult male and female mice. We found that Nrp2 is enriched in adult layer V pyramidal neurons, corticostriatal terminals, and in developing and adult striatal spiny projection neurons (SPNs). Loss of *Nrp2* increases SPN excitability and spine number, reduces short-term facilitation at corticostriatal synapses, and impairs goal-directed learning in an instrumental task. Acute deletion of *Nrp2* selectively in adult layer V cortical neurons produces a similar increase in the number of dendritic spines and presynaptic modifications at the corticostriatal synapse in the *Nrp2^-/-^* mouse, but does not affect the intrinsic excitability of SPNs. Furthermore conditional loss of *Nrp2* impairs sensorimotor learning on the accelerating rotarod without affecting goal-directed instrumental learning. Collectively, our results identify Nrp2 signaling as essential for the development and maintenance of the corticostriatal pathway and may shed novel insights on neurodevelopmental disorders linked to the corticostriatal pathway and semaphorin signaling.

**Significance Statement:** The corticostriatal pathway controls sensorimotor, learning and action control behaviors and its dysregulation is linked to neurodevelopmental disorders, such as autism spectrum disorder (ASD). Here we demonstrate that neuropilin 2 (Nrp2), a receptor for the axon-guidance cue semaphorin 3F, has important and previously unappreciated functions in the development and adult maintenance of dendritic spines on striatal spiny projection neurons (SPNs), corticostriatal short-term plasticity, intrinsic physiological properties of SPNs and learning in mice. Our findings, coupled with Nrp2’s association with ASD in human populations, suggest that Nrp2 may play an important role in ASD pathophysiology. Overall, our work demonstrates Nrp2 as a key regulator of corticostriatal development, maintenance and function, and may lead to better understanding of neurodevelopmental disease mechanisms.

## Introduction

The striatum constitutes the main input structure of the basal ganglia (BG). It receives massive excitatory projections originating from essentially all regions of the cortex as well as multiple thalamic nuclei (Buchwald, 1973; Kemp and Powell, 1970; (Smith, 2014). A major target of cortical input to the striatum is the dendritic spines of the GABAergic spiny projection neurons (SPNs), which constitute about 95% of striatal neurons (Gerfen and Surmeier, 2011). The corticostriatal pathway mostly originates from layer V of the cortex and to a lesser extent from layer III (McGeorge and Faull, 1989; Shepherd, 2013; Sohur et al., 2014). Corticostriatal projections in the adult animal are essential for the regulation of a wide range of behavioral and cognitive functions, and dysregulation of corticostriatal activity has been implicated in many neurodevelopmental and neuropsychiatric disorders (Balleine and O’Doherty, 2010; Pennartz et al., 2009; Shepherd, 2013; Shiflett and Balleine, 2011).

Many factors influence the organization and development of cortical projection neurons, including molecular guidance cues (Ding et al., 2012; Greig et al., 2013; Sohur et al., 2014). One such factor is the class 3 secreted semaphorins (Sema3s), a member of the semaphorin protein family that is essential for the appropriate elaboration of axons and dendrites during neural development (Mann et al., 2007; Pasterkamp and Giger, 2009; Tran et al., 2007; Yoshida, 2012). Sema3s utilize both Neuropilins (Nrp) and Plexins (Plxns) as their ligand binding and signal transducing receptor subunits, respectively, forming a receptor complex (Mann et al., 2007; Tran et al., 2007). In the developing mammalian nervous system, Sema3F operating through Nrp2/PlxnA3 ligand-receptor signaling has an important role in axon repulsion and in dendritic spine morphogenesis (Giger et al., 2000; Huber et al., 2005; Tran et al., 2009; Yaron et al., 2005). During mouse postnatal development, Sema3F regulates excitatory synapse formation in apical dendrites of layer V pyramidal neurons, and loss of this signaling increases apical dendrite spine density and the excitability of these neurons (Demyanenko et al., 2014; Tran et al., 2009). In humans, semaphorin and plexin gene variants are associated with a number of neurodevelopmental disorders, including autism spectrum disorder (Cheng et al., 2013; Fujii et al., 2011; Kim et al., 2017; Mosca-Boidron et al., 2016; Wu et al., 2007).

Semaphorins may have an important role in corticostriatal pathway development and maintenance in adult animals. Indeed, the related semaphorin Sema3E is present in the developing striatum and signals through PlexinD1 to establish thalamostriatal synapses on direct pathway SPNs (Ding et al., 2012). Furthermore, semaphorins have a recently described role in various forms of adult synaptic plasticity (Orr et al., 2017; Wang et al., 2017). Finally, semaphorin signaling could influence corticostriatal synapses by altering excitatory synapse formation in apical dendrites of Layer V neurons, which may have downstream effects on the corticostriatal synapse (Kozorovitskiy et al., 2012; Peixoto et al., 2016).

Here, we investigated the role of Sema3F-Nrp2 signaling in corticostriatal plasticity and related behaviors using either *Nrp2* null (knockout, KO) mice or mice in which *Nrp2* was conditionally deleted from adult layer V neurons (*Nrp2flox;Etv1-CreERT2* conditional knockout, CKO). In these mice, we analyzed the expression profile of Nrp2 in layer V cortical neurons and corticostriatal axons as well as in SPNs. Further, we tested whether global deletion of Nrp2 or conditional deletion in layer V cortical neurons altered corticostriatal short-term plasticity. Finally, we examined these mice in an instrumental learning task and the accelerating rotarod, behaviors known to depend on the corticostriatal pathway (Costa, 2007; Costa et al., 2004; Shiflett et al., 2015; Yin et al., 2009b). Altogether, our results demonstrate the importance of Sema3F-Nrp2 signaling in corticostriatal physiology and related behaviors.

## Materials and Methods

### Animals

The expression patterns and developmental phenotypes of the *Neuropilin 2* (*Nrp2*) knockout mice have been previously described in detail (Giger et al., 2000). The *Nrp2* floxed mice were previously described (Walz et al., 2002), where after Cre recombination part of exon 1 including the start codon is removed, and tauGFP-pA^+^ is placed immediately downstream of the Nrp2 promoter. The *Nrp2* floxed mice were crossed with the *Etv1-CreERT2* (Stock #013048; The Jackson Laboratory) mice where Cre recombinase was shown to be expressed specifically in layer V cortical neurons (Harris et al., 2014). Some *Nrp2f/f;Etv1+/Cre* progenies were crossed with the *Thy1-GFP* reporter (M line; Stock #007788, The Jackson Laboratory) to specifically detect GFP expression in cortical layer V pyramidal neuron (Feng et al., 2000). Mice used in this study were backcrossed for at least 10 generations to the C57BL/6NTac background strain. Only males were used for behavioral testing. During behavioral testing procedures, mice were placed on a restricted food diet of approximately 5 g of standard chow (Purina, St. Louis MO, USA) given in their home cage each day after behavior procedures were complete. Animals were weighed daily, and their body weights during the procedure were within 85-90% of their free-feeding weight. All procedures followed the NIH Guide to Care and Use of Laboratory Animals and were approved by the Rutgers Institutional Animal Care and Use Committee.

### Immunocytochemistry

Early postnatal P3, P14, P21 and adult (6-9 months) WT mice were subjected to transcardial perfusion with 4% paraformaldehyde (PFA) in phosphate buffered saline (PBS), removed and post fixed for 1 hour fixation with 4% PFA. For anti-Nrp2 labeling, tissue was prepared as previously described (Schneider Gasser et al., 2006). Briefly, brains from WT and *Nrp2^-/-^* mice were dissected and snap frozen using isopentane pre-chilled with dry ice. Brains were sectioned on a cryostat at 20µm along the coronal plane. Fresh frozen sections were fixed by brief immersion in 4% PFA for 2 minutes. Sections were processed for immunocytochemistry as previously described (Tran et al., 2013), with the following modifications. After PBS wash, the sections were incubated in 0.8% Triton X-100 in PBS for 10 minutes followed by 3 washes of 5 minutes each in PBS. Next, the sections were incubated in blocking solution (5% normal donkey serum, 0.1% Triton X-100, in PBS) for 1 hour. After blocking, the sections were incubated with primary antibodies overnight at 4**°** C. The next day, the sections were washed 3 times for 10 minutes each in PBS, followed by secondary antibody incubation for 2 hours in the dark. Finally, the sections were washed 4 times for 10 minutes each in PBS, slide mounted with Vectashield antifade mounting medium with Dapi (Vector Laboratories). Primary antibodies used were: rabbit monoclonal anti-Nrp2 (1:500, Cell Signaling Technology Cat# 3366, RRID:AB_2155250); goat polyclonal anti-Nrp2 (5µg/ml, R&D Systems); rabbit polyclonal anti-Darpp-32 (1:400, Cell Signaling Technology Cat# 2306, RRID:AB_823479); guinea pig polyclonal anti-Vglut1 (1:1000, Millipore); chicken polyclonal anti-GFP (1:1000, Aves Labs) and for visualization, AlexaFluor 488 or AlexaFluor 568 (1:500, Life Technologies); Cy5 (1:500, Jackson ImmunoResearch).

### Data analysis and photo-documentation

All immunocytochemically processed brain sections were analyzed using the Zen software (Zeiss) and Zeiss 510 LSC confocal system, and representative non-overlapping fields were imaged. Compiled Z stack images were exported as tif files. Images were captured with a Plan-Apochromat 63x/1.40 N.A. Oil immersion objective for high magnification, a Plan-Apochromat 20x/0.8 N.A. for lower magnification, and the following laser lines: 405nm, 488nm, 543nm, and 633nm.

### Preparation of brain slices for electrophysiology

Mice aged 3-6 months were deeply anesthetized with a mixture of ketamine/xylazine (80/20 mg/kg) and transcardially perfused with ice cold N-methyl D-glucamine (NMDG)-based solution containing (in mM): 103.0 NMDG, 2.5 KCl, 1.2 NaH_2_PO_4_, 30.0 NaHCO_3_, 20.0 HEPES, 25.0 Glucose, 101.0 HCl, 10.0 MgSO_4_, 2.0 Thiourea, 3.0 sodium pyruvate, 12.0 N-acetyl cysteine, 0.5 CaCl_2_ (saturated with 95% O_2_ and 5% CO_2_, pH 7.2-7.4). After decapitation, the brain was quickly removed and transferred into a beaker containing the ice-cold oxygenated NMDG-based solution and trimmed to a block containing the striatum. Coronal or oblique parahorizontal 300 µm sections were cut in the same medium using a Vibratome 3000 and immediately transferred to oxygenated NMDG-based solution at 35°C for 5 minutes, after which they were transferred to oxygenated normal Ringer’s solution at 25°C until placed in the recording chamber that was constantly perfused (2– 4 ml/min) with oxygenated Ringer’s solution at 32-34°C.

### Differential interference contrast imaging and recording

Slices were visualized under infrared–differential interference contrast microscopy with a digital frame transfer camera (Cooke SensiCam) mounted on an Olympus BX50-WI epifluorescence microscope with a 40X long working distance water-immersion lens.

Micropipettes for whole-cell recording were constructed from 1.2mm outer diameter borosilicate pipettes on a Narishige PP-83 vertical puller. The standard internal solution for whole-cell current-clamp recording was as follows (in mM): 130 K-gluconate, 10 KCl, 2 MgCl_2_, 10 HEPES, 4 Na_2_ATP, 0.4 Na_2_GTP, plus 0.1–0.3% biocytin, pH 7.3. These pipettes had a DC impedance of 3-4 MΩ. Membrane currents and potentials were recorded using an Axoclamp 700B amplifier (Molecular Devices). Recordings were digitized at 20 kHz with a CED Micro 1401 Mk II and a PC running Signal, version 5 (Cambridge Electronic Design). Sweeps were run at 20 s intervals.

Stimulating electrodes consisted of bipolar enamel/nylon-coated 100-µm-diameter stainless steel wires (California Fine Wire). For cortical stimulation, the electrodes were placed onto the surface of the corpus callosum. Stimuli consisted of single square-wave pulses (typically 0.01–1 mA, 0.3 ms duration). For paired-pulse ratio experiments, brief trains consisting of 2 pulses delivered at 5 to 50 Hz were generated.

### Biocytin histochemistry

At the completion of the experiments, slices containing biocytin-injected neurons were fixed by immersion in 4% PFA / 15% picric acid in PBS and stored overnight in this fixative solution at 4°C before washes in PBS. The sections were washed three times for 10 min each in 0.1 M phosphate buffer (PB) followed by 10% methanol and 3%H2O2 for 15 min After three washes for 10 min each in 0.1 M PBS, the sections were incubated with avidin–biotin peroxidase complex (Vector Laboratories; 1:200) and 0.1% Triton X-100 overnight at 4°C. After washing six times for 10 min each in 0.1 M PB, the sections were reacted with 3,3’-diaminobenzidine (DAB) (0.025%) and H_2_O_2_ (0.0008%) in PB. Nickel intensification was used (2.5mM nickel ammonium sulfate and 7 mM ammonium chloride in the DAB and H_2_O_2_ incubation). The sections were then post fixed in OsO_4_ (0.1% in PB) for 30 min, dehydrated through a graded series of ethanol, followed by propylene oxide, and infiltrated overnight with a mixture of propylene oxide and epoxy resin (Durcupan; Fluka Chemie). The sections were then transferred to fresh resin mixture for several hours and flat embedded between glass slides and coverslips and cured at 60°C for 24 hr.

### Dendritic spine quantifications

SPNs labeled with biocytin were photographed with a Zeiss Axiovert 200M inverted microscope, 40x objective, saved as compiled Z stack images and exported as tif files to Image J for quantification of individual spines. All spines located between 50 to 100 μm from the cell soma on three randomly selected dendrites of each SPN were counted and the average number of spines per 50 μm of dendritic length computed. This location was selected because the peak spine density in rodents occurs at about 60 μm from the soma (Wilson, 1993). At least 8-11 biocytin labeled SPNs per genotype were counted and n= 4 animals per genotype. Spine density (spines per μm) along *Thy1-GFP+* tagged layer V pyramidal neurons in adult brains 8-15 months old were imaged in the coronal plane and quantified using the following parameters: proximal to soma was defined as the first 45 μm. The number of spines was scored by counting spines along a 45 μm segment of the apical dendrite. The middle regions were defined as 45 – 90 μm from the soma. The number of spines was scored by counting spines along this 45μm segment of the apical dendrite. A total of 60 layer V pyramidal neurons were scored for each of *Nrp2^f//f^;Etv1-CreER+* and littermate controls; 3 animals were used per genotype. Littermate control genotypes include: *Nrp2^f//f^;Etv1^+/+^;Thy1-GFP+, Nrp2^+//f^;Etv1^+/+^;Thy1-GFP+ and Nrp2^+//f^;Etv1^+/Cre^;Thy1-GFP+*.

### Behavioral Procedures

#### Instrumental behavior

Mice were tested in four identical operant conditioning chambers (Med Associates, St. Albans VT, USA). Each operant conditioning chamber measured 15.9 cm x 14.0 cm x 12.7 cm (w/h/d) and was constructed of stainless steel and clear plastic walls and a stainless-steel grid floor. A food cup with infrared detectors was centered on one wall with retractable levers situated to the left and right of the food cup. Responses on one lever caused delivery of a single 20 mg grain-based chocolate flavored food pellet (Bio-serv, Frenchtown NJ USA) into the food cup from a dispenser mounted outside the operant conditioning chamber. A single stimulus light located above the lever illuminated for 2 sec during food pellet delivery. A 28 V light was located on the opposite wall from the food cup and illuminated the operant conditioning chamber during behavioral procedures. Each operant conditioning chamber was housed in a sound-attenuating shell and equipped with a ventilation fan that was activated during behavioral procedures. Control over the operant conditioning chambers was enabled by a PC operating through an interface. Operant conditioning chamber operation and data collection were carried out with Med Associates proprietary software (Med-PC IV).

Behavioral procedures commenced after one week of food restriction. Mice were habituated to the operant conditioning chamber in one 15-min session, which was followed the next day by a 20 min session in which food pellets were dispensed on a random-time (RT) 60 sec schedule accompanied by 2-sec illumination of the stimulus light. The levers were withdrawn during this phase. The following day mice began daily instrumental training sessions. During each session, a single lever was inserted into the operant conditioning chamber and responses on the lever delivered a food pellet. For the first 2 sessions each response resulted in pellet delivery and illumination of the stimulus light. For the remaining 8 sessions, outcomes were delivered according to a variable-ratio (VR) schedule, which required, on average, 7.5 responses to issue pellet delivery and illuminate the stimulus light. We used this training schedule because it has been shown to maintain goal-directed responding (Adams and Dickinson, 1981)

Following instrumental conditioning, mice underwent a selective-satiety induced outcome devaluation test. Mice were placed in individual cages identical to their home cages with a bowl that contained 10 g of food pellets. The bowl contained either the same flavor of food pellets as the instrumental outcome or an alternative flavor. After 1 hour, mice were placed in the operant conditioning chamber and the lever was inserted for a 10-min extinction test. No pellets or stimulus lights were delivered during the test. The test was repeated the following day with the opposite type of food pellet from the first test serving as the devalued outcome.

#### Open Field

The open field apparatus measured 40 cm x 40 cm and was constructed of opaque gray plastic with a solid metal base (Stoelting Co, USA). The apparatus had overhead lighting and the illumination at the center of the base was approximately 400 lumens. Mice were placed in the center of the open field and removed after 20 minutes. The test was conducted twice, once per day for two consecutive days. The data portrayed in Supplemental Figure S4 represents the average of two sessions. A camera (Basler Ace 60 fps) mounted above the open field was used to record behavior during the test session. The distance traveled and time spent in center zone (20 cm x 20 cm) during the test was calculated using tracking software (Noldus ethovision v. 11.5).

#### Elevated Zero Maze

The elevated zero maze (Stoelting Co) is a circular elevated track 50 cm in diameter. The track lane is 5 cm in width. The track is raised 50 cm from the floor. One half of the track is enclosed in walls that are 15 cm in height. The track and walls are constructed of opaque material. During the test trial, a mouse was placed in an enclosed portion of the maze and allowed to freely explore. After 5 minutes the mice were removed. If mice fell from the maze in the first two minutes of the trial then the trial was repeated, otherwise the trial was aborted. A video camera mounted above the maze captured the mouse’s behavior during the trial. Noldus ethovision was used to track the mouse’s position during the trial.

#### Accelerating Rotarod

The single-lane rotarod (Med Associates, St. Albans VT) consisted of a 3.18 cm diameter grooved plastic spindle raised 30 cm from the base of the apparatus. Mice were placed on the spindle, which linearly accelerated from 4 to 40 rpm over 2 min. The trial ended when the mouse fell off the spindle, made one complete revolution while gripping the spindle, or 5 min had elapsed. Each mouse received 5 consecutive trials per day for two days.

## Results

### Expression profiles and effects of Neuropilin 2 on the corticostriatal pathway

We previously showed that the ligand Sema3F is expressed in the postnatal cortex, while the receptor Nrp2 is uniquely localized to the apical dendrites of cortical neurons *in vitro* (Tran et al., 2009). First, we asked whether this specific localization of Nrp2 in apical dendrites could be recapitulated *in vivo* in layer V pyramidal neurons that are known to project to striatal neurons. We used a previously validated anti-Nrp2 antibody (Tran et al., 2013), and demonstrated that Nrp2 is localized in layer V cortical neuron soma and apical dendrites from adult (6-9 months old) WT, but not in *Nrp2* global KO mice (Fig. 1A-D). Interestingly, very little or no Nrp2 is detected in basal dendrites, which is also consistent with our previous *in vitro* findings. Further, we performed a triple immunofluorescent staining using anti-DARPP32 (Ouimet et al., 1984), anti-Nrp2 and anti-VGLUT1 antibodies to detect the presence of Nrp2 in several VGLUT1-positive corticostriatal terminals in close apposition to DARPP32-positive processes in the striatum of WT adult brain sections but not in Nrp2 null brain sections (Fig. 1E, see merged panels and insets). This suggests that Nrp2 is present at the level of corticostriatal presynaptic axon terminals, where it may play a role in the development of axospinous corticostriatal synapses and the maintenance of synaptic activity in SPNs. Next, using double immunofluorescent staining with anti-Nrp2 and anti-DARPP-32, we detected co-localization of Nrp2 and DARPP32 within the same SPNs as early as postnatal day (P)3, through postnatal at P14, P21, and also in adult (6-9 months) SPNs (Fig. 1F). Nrp2 expression in SPNs seems to be present in both the somatic compartment and the dendritic processes (Fig. 1F).

**Figure 1.**
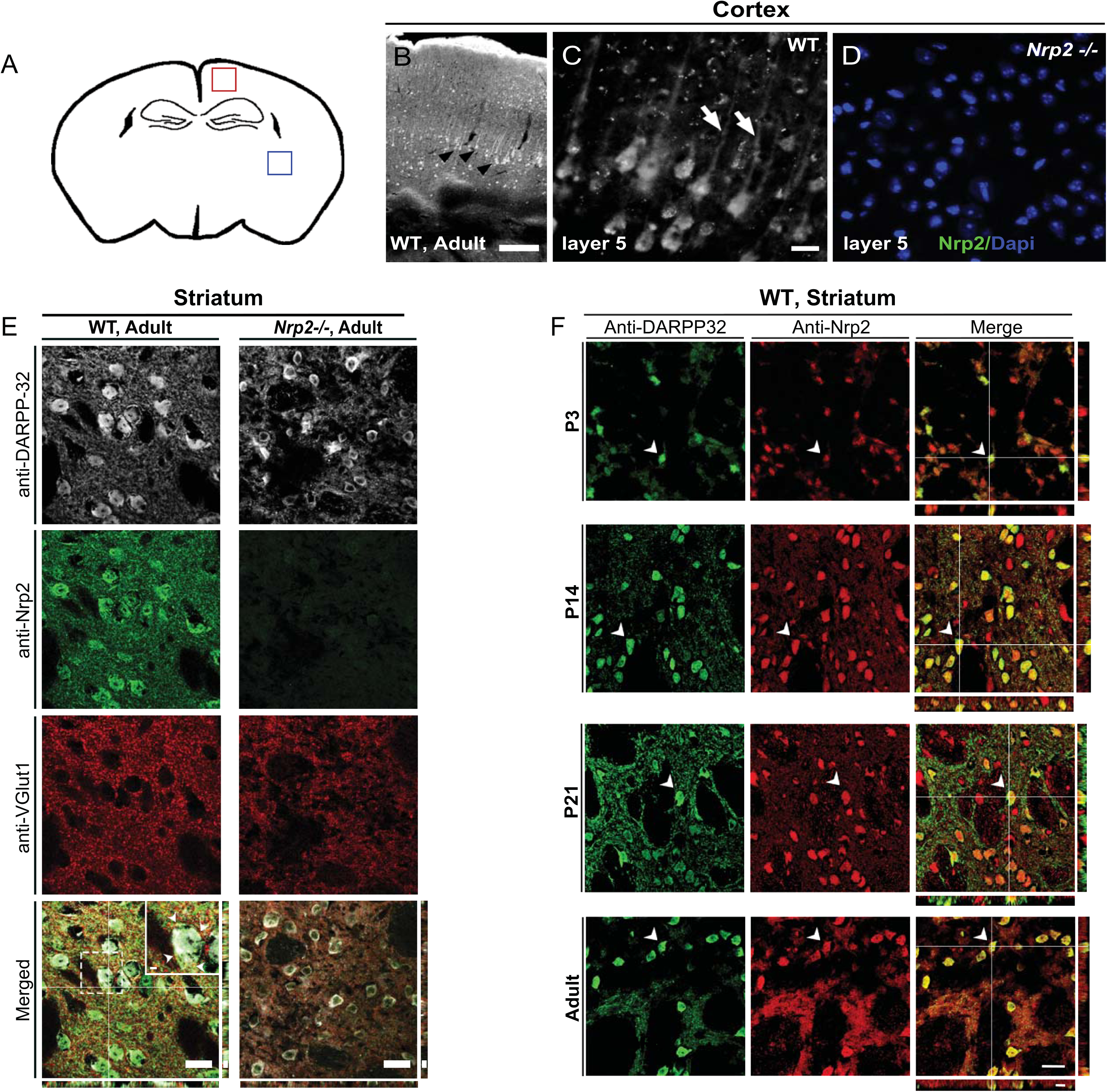
Nrp2 protein is expressed in cortical neurons and striatal SPNs. **A)** Schematic of a coronal brain section where representative confocal images were obtained for imaging the cortex (red box) and striatum (blue box). **B)** Expression of Nrp2 is detected in cortical neurons (black arrows), mainly in layer V, and sparsely in layers 2/3 and 6. **C)** High magnification of Layer V cortical neurons reveals Nrp2 localization in pyramidal neuron soma and apical dendrites. **D)** No expression is observed in adult *Nrp2^-/-^* cortical neurons. **E)** Adult WT (left column) and *Nrp2^-/-^* (right column) sections triple labeled with anti-Nrp2, anti-VGLUT1, and anti-Darpp-32. Nrp2 expression is detected in VGLUT1^+^ terminals (in Merged panel, white arrowheads, boxed inset). **F)** WT P3, P14, P21 and adult brain sections co-immunolabeled with anti-Nrp2 (red) and anti-Darpp-32 (green). SPNs co-expressing Nrp2 and Darpp-32 (white arrowheads). Scale bar 200μm in B; 50μm in C for C and D; 20μm in all panels for E, except 8μm in Z-plane; and 20μm in all panels for F, except 8µm in Z-plane.

### Modifications in intrinsic electrophysiological properties of SPNs in Nrp2^-/-^ mutant mice

Next, we investigated whether the loss of Nrp2 signaling affects the intrinsic electrophysiological properties of SPNs. We tested responses to injected current pulses in SPNs recorded in whole-cell current clamp from WT and *Nrp2^-/-^* animals. We found a significant increase in the depolarization to current injection at higher current steps in *Nrp2^-/-^* mice compared to WT controls (Fig. 2A-C). Also, spike frequency in response to depolarizing somatic current injection was significantly increased in SPNs from *Nrp2^-/-^* mice compared to WT controls (Fig. 2D). Consistent with an increased intrinsic excitability of SPNs in *Nrp2^-/-^*mice, the rheobase current was significantly lower in the KO mice compared to WT SPNs. (366.41 ± 13.93 vs. 315.69 ± 18.39 for WT and *Nrp2^-/-^* respectively; Fig. 2E-F; *p* = 0.011, unpaired t-test, 13-20 neurons/animal/genotype, averaged from n=4 animals/genotype). Analysis on the action potential waveform revealed no significant differences between the WT and *Nrp2^-/-^* mice (data not shown).

**Figure 2.**
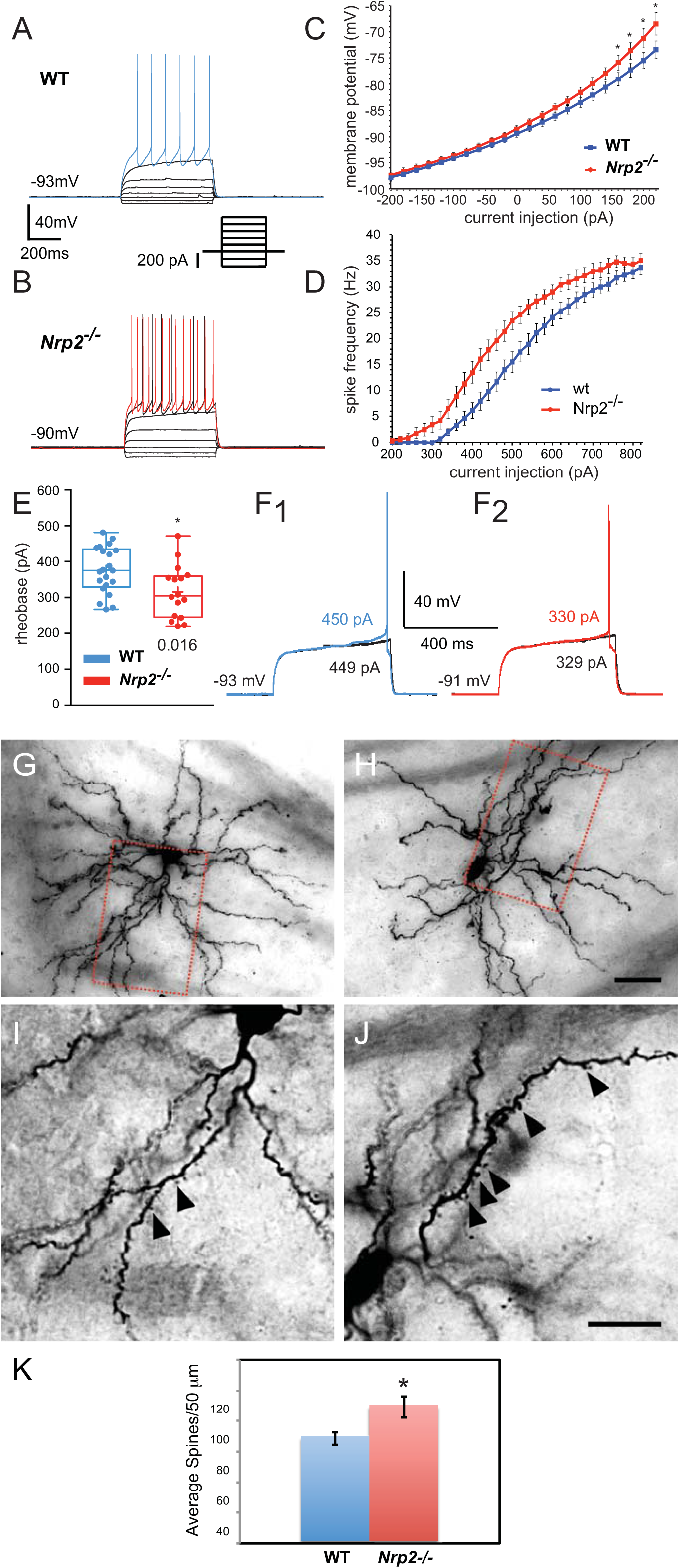
Changes in SPNs intrinsic properties and spine density in *Nrp2^-/-^*mice. **A-B)** Membrane voltage responses to injected current pulses of a SPN recorded in a WT (A) and *Nrp2^-/-^*(B) mice. Note that similar current injection elicits higher firing frequency in *Nrp2-/-* mice (B, red) than in WT mice (A, blue). **C)** Current voltage curves of SPNs recorded in WT (blue) and *Nrp2^-/-^*mice (red). **D)** Graph representing the spike frequency in response to injected current pulses in WT (blue) and *Nrp2^-/-^* mice (red). **E)** Box plots representing the rheobase current in WT (blue) and *Nrp2^-/-^* mice (red). **F)** Representative current clamp traces showing the rheobase current of a SPN recorded in WT (F1) and *Nrp2^-/-^* mice (F2). Note that the rheobase current is lower in F2 (*Nrp2^-/-^*) than in F1 (WT). **G-H)** Representative images taken from a WT and *Nrp2^-/-^* SPN back-filled with biocytin. 2-D confocal images are reconstructed from complete z-stacks. **I-J)** Single-plane images of WT and *Nrp2^-/-^* SPNs were taken at higher magnification within the area indicated by the red-dashed boxes in G and H, respectively. **K)** Quantification of average number of spines per 50 μm from reconstructed z-stack images of 2-D neurons. Error bars are ± SD, unpaired t-test, \**p* = 0.0022. Scale bars H=20 μm for G and H; J=20 μm for I and J.

### *Nrp2^-/-^* mice exhibit supernumerary spines in adult spiny projection neurons

Our previous findings in layer V pyramidal neurons in the cortex showed that Nrp2 plays a role to restrain spine number (Tran et al., 2009). Here, we examined whether Nrp2 has a similar effect in SPNs by measuring spine density in biocytin-stained SPNs following whole-cell recordings in *Nrp2^-/-^* and control mice. While we did not observe any gross dendritic morphological changes (Fig. 2G-H), we did find on average a ∼23% increase in spine density on dendrites between 50 to 100 μm away from the cell soma in *Nrp2^-/-^* compared to WT SPNs (Fig. 2I-K; 68.63 ± 4.17 vs. 89.18 ± 6.87 for WT and *Nrp2^-/-^* respectively; unpaired *t*-test *t*_6_ = 5.116, *p* = 0.0022). These results suggest that Nrp2 play an important role in regulating spine density in SPNs.

### *Nrp2^-/-^* mice exhibit impaired paired-pulse facilitation from cortical-stimulated input to the striatum

Next, we asked whether loss of Nrp2 altered cortical excitatory input to SPN’s. To address this, we stimulated corticostriatal inputs while recording striatal SPNs in *Nrp2^-/-^* and wild type (WT) adult mice (Fig. 3A-B). Short-term plasticity was tested by using double pulse stimulation with a range of different interstimulus intervals (ISI, 20, 40, 60, 120, 200 ms) and we calculated the paired pulse ratio (PPR). As previously described, corticostriatal synapses measured in WT SPNs exhibit paired pulse facilitation indicating a low release probability (Ding et al., 2008). Here, the PPR was significantly reduced in SPNs from *Nrp2^-/-^* neurons in comparison to WT (PPR_20msWT_ = 1.764 ± 0.11 vs PPR_20msCKO_ = 1.19 ± 0.08; *p*=0.0009; PPR_40msWT_ = 1.69 ± 0.13 vs PPR_40msCKO_ = 1.11 ± 0.07; *p*=0.001; PPR_60msWT_ = 1.61 ± 0.13 vs PPR_60msCKO_ = 0.99 ± 0.05; *p*=0.0002; PPR_80msWT_ = 1.47 ± 0.12 vs PPR_80msCKO_ = 1.06 ± 0.09; p=0.01; PPR_100msWT_ = 1.36 ± 0.10 vs PPR_100msCKO_ = 1.05 ± 0.09; *p*=0.03; PPR_120msWT_ = 1.36 ± 0.13 vs PPR_120msCKO_ = 0.98 ± 0.04; *p*=0.007; PPR_200msWT_ = 1.21 ± 0.07 vs PPR_200msCKO_ = 0.97 ± 0.07; *p*=0.03; n=14-22 SPNs per condition; Fig. 3C-F). Grouped analysis also reveals significant differences between the *Nrp2*^-/-^ mice and their littermate controls for corticostriatal short-term plasticity (Fig. 3G; F_1, 213_ = 62.41; *p* = 0.0001, two-way ANOVA, Fig. 3G). Altogether, these observations indicate that loss of Nrp2 significantly increases initial release probability from corticostriatal synapses.

**Figure 3.**
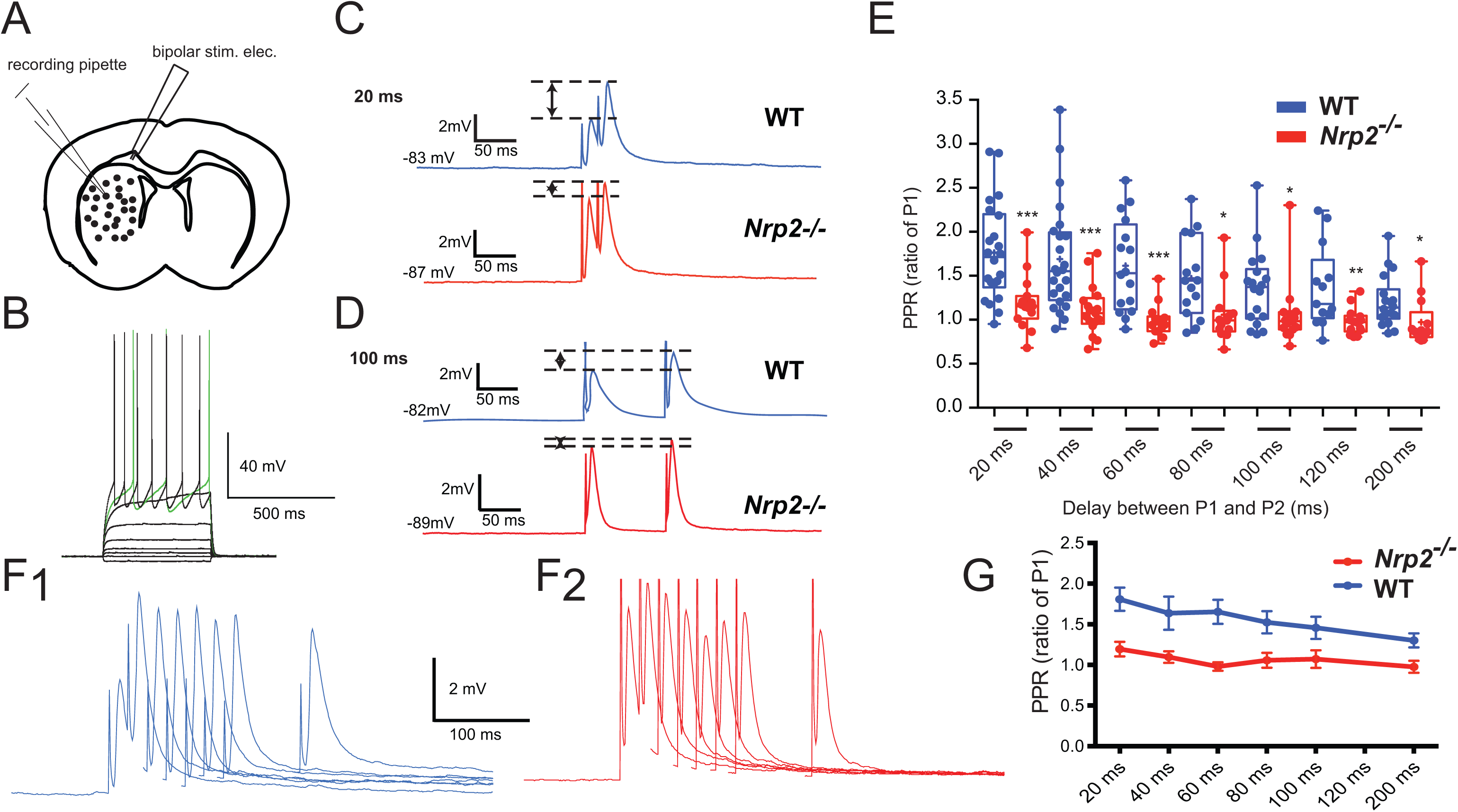
Defects in corticostriatal short-term plasticity onto SPNs in *Nrp2^-/-^* mice. **A)** Schematic illustrating the experimental paradigm where SPNs where recorded in coronal striatal slices. A bipolar stimulating electrode was placed in the corpus callosum to stimulate corticostriatal fibers. **B)** Example of voltage responses to current steps injection of a SPN. **C-D)** Representative examples of EPSPs measured in SPNs after a pair of electrical stimuli. Interstimulus interval (ISI) 20 ms in C and 100 ms in D. Responses of SPNs recorded in WT are represented in blue and mutant in red. Note that in WT SPNs there is a strong paired pulse facilitation that is impaired in *Nrp2^-/-^.* **E)** Box plots representing the paired-pulse ratio of the responses at different ISI (20, 40, 60, 80, 100, 120 and 200ms) for the 2 genotypes. Responses are expressed as a ratio of the response to the first stimulus. **F)** Representative current clamp traces showing corticostriatal EPSPs elicited by paired stimuli with increasing ISIs in WT mice (F1) and *Nrp2^-/-^*(F2). G Summary graph of PPRs recorded from medium spiny neurons plotted against ISI for cortical stimulation in WT (blue) and *Nrp2^-/-^* (red). Box plots represent the minimum, maximum interquartile range, the mean and median. Statistical analysis was made using unpaired t-test (E) and two-way ANOVA (G).

### *Nrp2^-/-^* mice are impaired in goal-directed instrumental behavior

Based on the observed structural and physiological alterations in cortical and striatal neurons in *Nrp2^-/-^*mice, we examined performance in an operant task known to depend on the corticostriatal pathway (Hart et al., 2014; Shiflett and Balleine, 2011). We trained mice to make lever press responses to obtain food pellets. All mice learned the task, and response rates increased over the course of training, as shown by a significant main effect of training session on response rate (repeated measures ANOVA: F_9, 144_ = 13.97, *p* = 0.0001) (Fig. 4A). Overall, *Nrp2^-/-^*mice responded at a significantly lower rate compared to WT mice, as shown by a main effect of genotype on response rate (ANOVA: F_1, 16_ = 5.50, *p* = 0.036). Furthermore, *Nrp2^-/-^* and WT mice differed in the rate of acquisition of the task. By the 10^th^ training session, WT mice responded at a significantly greater rate compared to *Nrp2^-/-^* mice, but no significant difference was observed on sessions 1-9 (Fig. 4B).

**Figure 4.**
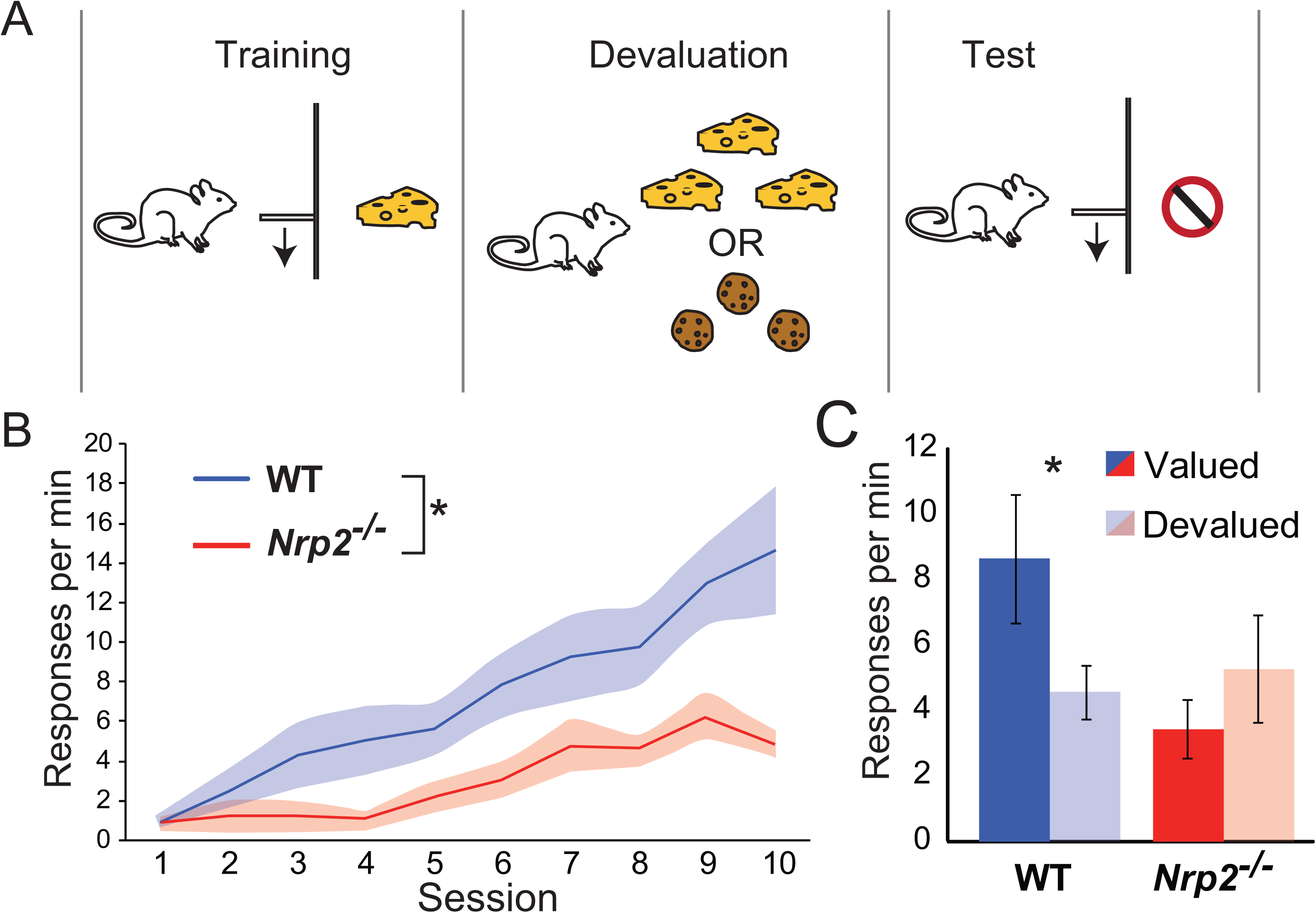
Mice carrying a deletion mutation to *Nrp2* are impaired in goal-directed instrumental action. **A)** We trained *Nrp2^-/-^* and WT mice on an instrumental learning task with a selective satiety procedure to devalue the instrumental outcome or an alternative outcome. **B)** All mice acquired an instrumental response that increased in rate with training; however, the rate of acquisition was reduced in *Nrp2^-/-^*mice. **C)** When the instrumental outcome was devalued, WT mice significantly reduced their responding compared to when an alternative food was devalued. In contrast, *Nrp2^-/-^* mice responded at similar rates following both devaluation sessions. Two-way ANOVA, * = *p* < 0.05; error bars = ± 1 SEM.

We next examined goal-directed behavior in *Nrp2^-/-^*mice. Goal-directed behavior is evident when animals flexibly alter their actions in response to changes in outcome value (Adams and Dickinson, 1981). This behavioral flexibility is dependent on corticostriatal pathway plasticity (Hawes et al., 2015; Shan et al., 2014; Shiflett et al., 2010). To test goal-directed behavior, we used a selective-satiety procedure to reduce the value of the instrumental outcome and then probed the animals’ responses in an extinction paradigm. This test occurred after animals had acquired an instrumental response. Feeding WT mice the pellets used during instrumental training significantly reduced their subsequent responding during the extinction test, compared to responses they made after being fed an alternative flavor of pellets (paired *t-*test, *t*_10_ = 2.27, *p* = 0.046) (Fig. 4C). In contrast, *Nrp2^-/-^* mice responded at similar rates during the extinction tests, whether following feeding on the instrumental outcome or an alternative flavor of pellets (Fig. 4C). Overall, we observed a significant outcome value x genotype interaction (ANOVA: F_1, 16_ = 4.95, *p* = 0.041) on response rates during the devaluation tests. The lack of sensitivity to changes in outcome value in *Nrp2^-/-^ mice* was not a consequence of differences in food consumption. We found no difference between genotypes on the amount of food pellets consumed during the satiety procedure (data not shown). This suggests that the inability to adjust responding following changes in the desirability of an outcome likely reflects impaired acquisition or control of goal-directed behavior.

### Acute deletion of Nrp2 in adult layer V cortical neurons increased apical dendritic spine number

We addressed the question of whether the different alterations measured in the global *Nrp2^-/-^* KO animals could be attributed to Nrp2 expression in cortical layer V neurons or striatal neurons (See Fig. 1). We used the conditional floxed *Nrp2* mouse line (Walz et al., 2002) crossed to the Cre recombinase-ERT2 driver line under the regulation of the Etv1 promoter, which was previously demonstrated to be expressed specifically in layer V cortical neurons (Harris et al., 2014; Fig. 5A and 5B). The *Nrp2* conditional knockout (CKO) cassette contains a tauGFP-pA^+^ insert that will be expressed under the *Nrp2* promoter upon successful Cre recombination. After generating the *Nrp2f/f;Etv1-CreERT2* animals and treating them with tamoxifen at 4 to 6 months of age, we confirmed specific deletion of Nrp2 in layer V cortical neurons by performing immunofluorescent staining with anti-GFP primary antibodies on *Nrp2f/f;Etv1+/+* (control) or *Nrp2f/f;Etv1+/Cre* (CKO) mouse brain sections. We detected GFP-positive cortical pyramidal neurons with apical dendrites that extended to the pial surface in adult *Nrp2f/f;Etv1+/Cre* CKO mice treated with tamoxifen but not in *Nrp2f/f;Etv1+/+* control littermates (Fig. 5C-G). Importantly, we found no expression of GFP-positive SPNs in *Nrp2f/f;Etv1+/Cre* CKO or *Nrp2f/f;Etv1+/+* mouse control brains further demonstrating the specificity of our acute Nrp2 deletion in adult layer V cortical neurons (Fig. 5C, 5H-K).

**Figure 5.**
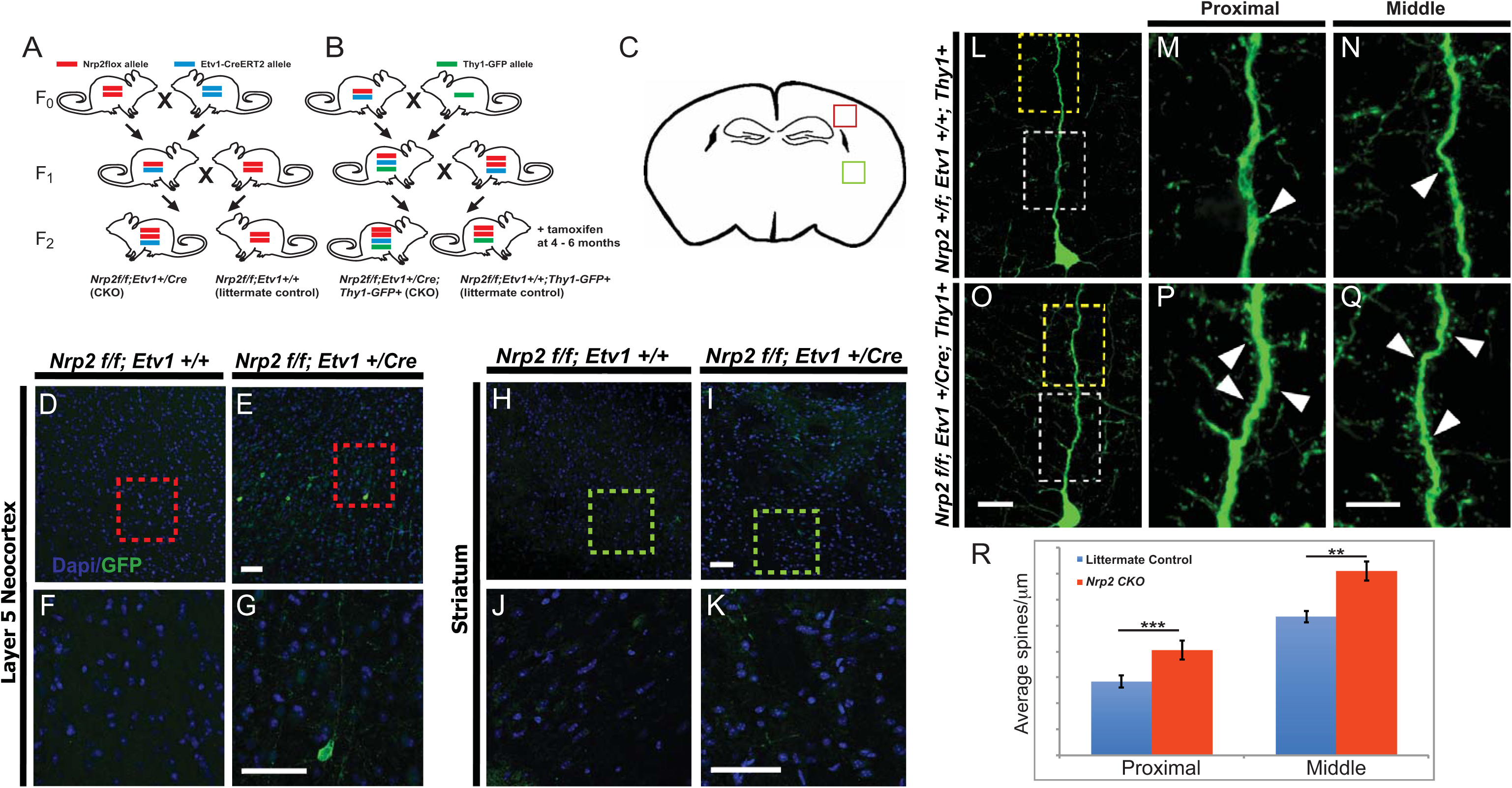
Specific deletion of *Nrp2* in adult layer V pyramidal neurons leads to increased dendritic spine numbers *in vivo*. **A**) *Nrp2flox* mice (F_0_) were crossed with *Etv1-CreERT2* (F_0_) mice to generate heterozygous (F_1_), the F_1_ were backcrossed with *Nrp2flox* mice to generate homozygous floxed *Nrp2flox* mice with or without *Etv1-Cre* positive alleles (F_2_). **B**) F_1_ Progeny developed in A were crossed with *Thy1-GFP* positive mice to generate triple heterozygous *Nrp2+/f;Etv1+/Cre;Thy1-GFP+*. Triple heterozygous were crossed with *Nrp2f/f;Etv1+/Cre* to generate *Nrp2f/f;Etv1+/Cre;Thy1-GFP+* and littermate controls (F_2_). All F_2_ in both A and B were treated twice on consecutive days with tamoxifen at 4-6 months. **C-K)** Schematic of a coronal adult brain section, red box indicates cortical region of image in (D-G) immune-labeled with anti-GFP showing successful Cre recombination in layer V neurons (E, red box enlarged in G) but not in littermate control (D, red box enlarged in F). Green box indicates striatal region of image in (H-K), no GFP-positive striatal neuron was detected in H (green box enlarged in J) or in I (green box enlarged in K). Scale bars, 50μm in E and I for D, E, and H, I, respectively; 50μm in G and K for F, G and J, K, respectively. **L-N)** Representative image of a neuron from a *Nrp2+/f;Etv1+/+;Thy1-GFP+* adult mouse brain. **O-Q)** Representative image of a neuron from a *Nrp2f/f;Etv1+/Cre;Thy1-GFP+*Thy1 adult mouse brain. Yellow and white boxes outline middle regions, 45-90μm from the soma and proximal regions, 0-45μm from the soma, respectively. White arrows show dendritic spines in proximal regions, M and P, enlarged from yellow box region in L and O, respectively. White arrows show dendritic spines in middle regions, N and Q, enlarged from yellow box region in L and O, respectively. **R)** Quantification of spine density on layer V apical dendrites in proximal and middle regions. A significantly higher more spines are found on *Nrp2f/f;Etv1+/Cre;Thy1-GFP+* neurons compared to littermate controls for both regions. Error bars are ± SEM; unpaired *t*-test; \*\**p*=0.0003; \*\*\**p*<0.0001 compared to littermate controls. Scale bars, 15μm in O for L and O, 15μm in Q for M, N, P and Q.

In order to determine whether Nrp2 in adult layer V neurons is required for the maintenance of apical dendritic spine number, as observed in the *Nrp2* null, we crossed *Nrp2f/f;Etv1+/Cre* and *Nrp2f/f;Etv1+/+* mice with the Thy1-GFP M-reporter line, where GFP is expressed sparsely in layer V pyramidal neurons (Fig 5B and 5C). We found that *Nrp2f/f;Etv1+/Cre;Thy1-GFP+* (CKO) animals treated with tamoxifen at 4 to 6 months of age exhibited a significantly increased number of apical dendritic spines (Fig. 5L-Q), ∼1.5-fold more in the proximal (0-45 μm from soma, 0.41 ± 0.02 spines/μm in CKO vs. 0.28 ± 0.02 spines/μm in control mice, *t_118_* = 4.078, *p* < 0.0001, unpaired t-test; Fig. 5R) and ∼1.3-fold more in the middle dendritic regions (45-90 μm from soma, 0.71 ± 0.04 spines/μm in CKO vs. 0.53 ± 0.03 spines/μm in control mice, *t_118_* = 3.775, *p* = 0.0003 unpaired t-test; Fig. 5R). This finding suggests that Nrp2 plays a role in dendritic spine maintenance in adult layer V cortical neurons.

### Loss of Nrp2 in adult cortical layer V projection neurons alters corticostriatal short-term plasticity but preserves intrinsic electrophysiological properties in SPNs

To address whether specific deletion of Nrp2 in adult layer V cortical neurons influences corticostriatal short-term plasticity and/or affects SPNs intrinsic physiological properties, we performed whole-cell recordings of SPNs in both *Nrp2f/f;Etv1+/Cre* (*Nrp2* CKO) and *Nrp2f/f;Etv1+/+* (WT littermate control) brain slices from adult mice treated with tamoxifen at 4 to 6 months of age. In contrast with results obtained using *Nrp2^-/-^*mice, SPNs recorded from *Nrp2* CKO mice show no significant changes in intrinsic excitability in comparison with WT littermates. Indeed, we measured similar current-voltage responses (Fig. 6A-C, n=16 WT, n=15 CKO), spike frequency (Fig. 6D, n=14 WT; n=14 CKO), input resistance (Fig. 6E, n=17 WT; n=15 CKO), resting membrane potential (Fig. 6F, n=17 WT; n=15 CKO) as well as in the rheobase current of SPNs (Fig. 6G-I, n=17 WT; n=14 CKO). These results strongly suggest that the changes in intrinsic excitability of SPNs we observed in the full *Nrp2^-/-^* mice are due to Nrp2 expression in striatal SPNs themselves and not in deep-layer cortical neurons.

**Figure 6.**
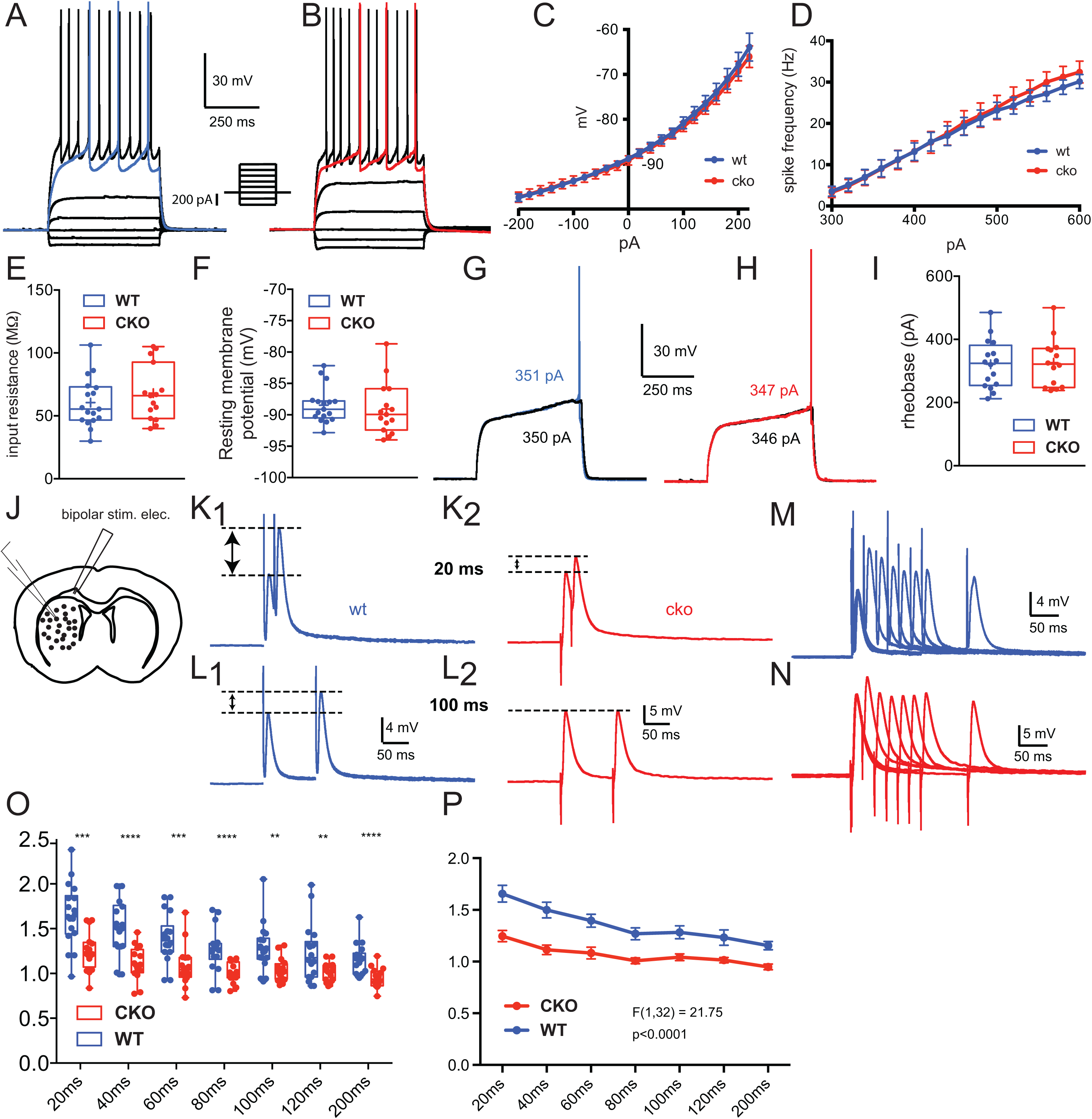
SPNs intrinsic properties and corticostriatal short-term plasticity in *Nrp2* CKO mice. **A-B)** Membrane voltage responses to injected current pulses of a SPN recorded in a WT (A) and CKO *Nrp2* (B) mice. **C)** Current voltage curves of SPNs recorded in WT (blue line) and CKO *Nrp2* mice (red line). **D)** Graph representing the spike frequency in response to injected current pulses in WT (blue line) and CKO *Nrp2* mice (red line). **E)** Box plots representing the input resistance measured at rest (WT: blue; CKO *Nrp2*: red). **F)** Box plots representing the resting membrane potential (WT: blue; CKO *Nrp2*: red). **G-H)** Representative current clamp traces showing the rheobase current of a SPN recorded in WT (G) and CKO *Nrp2* mice (H). **I)** Box plots representing rheobase current (WT: blue; CKO *Nrp2*: red). **J)** Schematic illustrating the experimental paradigm where SPNs where recorded in coronal striatal slices. A bipolar stimulating electrode was placed in the corpus callosum to stimulate corticostriatal fibers. **K-L)** Representative examples of EPSPs measured in SPNs after a pair of electrical stimuli. Interstimulus interval (ISI) 20 ms in K and 100 ms in L. (K1, L1: WT; K2 and L2: CKO *Nrp2* mice). Note that in WT SPNs there is a strong paired pulse facilitation that is impaired in CKO *Nrp2*. **M-N)** Representative current clamp traces showing corticostriatal EPSPs elicited by paired stimuli with increasing ISIs in WT mice (M) and CKO *Nrp2* (N). **O)** Box plots representing the paired-pulse ratio of the responses at different ISI (20, 40, 60, 80, 100, 120 and 200ms) for the 2 genotypes. Responses are expressed as the amplitude of the 2nd EPSP divided by the amplitude of 1st one. **P)** Summary graph of PPRs recorded from medium spiny neurons plotted against ISI for cortical stimulation in WT (blue) and CKO *Nrp2* (red). Box plots represent the minimum, maximum interquartile range, the mean and median. Statistical analysis was made using unpaired t-test (F) and two-way ANOVA (G).

Next, we tested the short-term plasticity at corticostriatal synapses in the *Nrp2* CKO mice in comparison with WT littermates. As above, we used electrical cortical stimulation with a range of different interstimulus intervals (ISI, 20, 40, 60, 120, 200 ms). Here, similar to the results obtained in *Nrp2^-/-^* mice, we measured a strong alteration in the PPR at corticostriatal synapses in *Nrp2* CKO (Fig. 6J-P; PPR_20msWT_ = 1.674 ± 0.091 vs PPR_20msCKO_ = 1.248 ± 0.0505; *p*=0.0003; PPR_40msWT_ = 1.516 ± 0.072 vs PPR_40msCKO_ = 1.124 ± 0.044; *p*<0.0001; PPR_60msWT_ = 1.416 ± 0.054 vs PPR_60msCKO_ = 1.091 ± 0.0538; *p*=0.0002; PPR_80msWT_ = 1.301 ± 0.061 vs PPR_80msCKO_ = 1.01 ± 0.026; *p*<0.0001; PPR_100msWT_ = 1.294 ± 0.064 vs PPR_100msCKO_ = 1.048 ± 0.031; *p*=0.0016; PPR_120msWT_ = 1.268 ± 0.078 vs PPR_120msCKO_ = 1.013 ± 0.02627; *p*=0.0043; PPR_200msWT_ = 1.171 ± 0.042 vs PPR_200msCKO_ = 0.949 ± 0.0258; *p*<0.0001; n=15-18 SPNs per condition). Grouped analysis also reveals significant differences between the littermate controls and *Nrp2* CKO animals for corticostriatal short-term plasticity (Fig. 6P; F_1, 32_ = 21.75; *p*<0.0001, two-way ANOVA).

Altogether, these results strongly support the hypothesis that acute deletion of Nrp2 specifically in layer V cortical neurons is sufficient to alter corticostriatal short-term plasticity, replicating the data obtained in the global *Nrp2^-/-^* mice.

### Loss of Nrp2 in adult deep layer cortical neurons selectively impairs sensorimotor learning

Next, we asked whether acute deletion of Nrp2 in adult layer V cortical neurons affects motor learning, anxiety, and goal-directed behaviors. We tested adult male *Nrp2* CKO and littermate control mice on the open field test, the elevated zero maze, the accelerating rotarod, and instrumental behavior. We found no differences between genotypes on locomotor behavior as measured by distance traveled in the open field test (Fig. 7A and 7B), nor any effect on time spent in the center zone of the open field, suggesting no effect on anxiety (Fig. 7B). We confirmed this result in the elevated zero maze test, which also showed no effect of genotype on time spent in the open arms (Fig. 7C).

**Figure 7.**
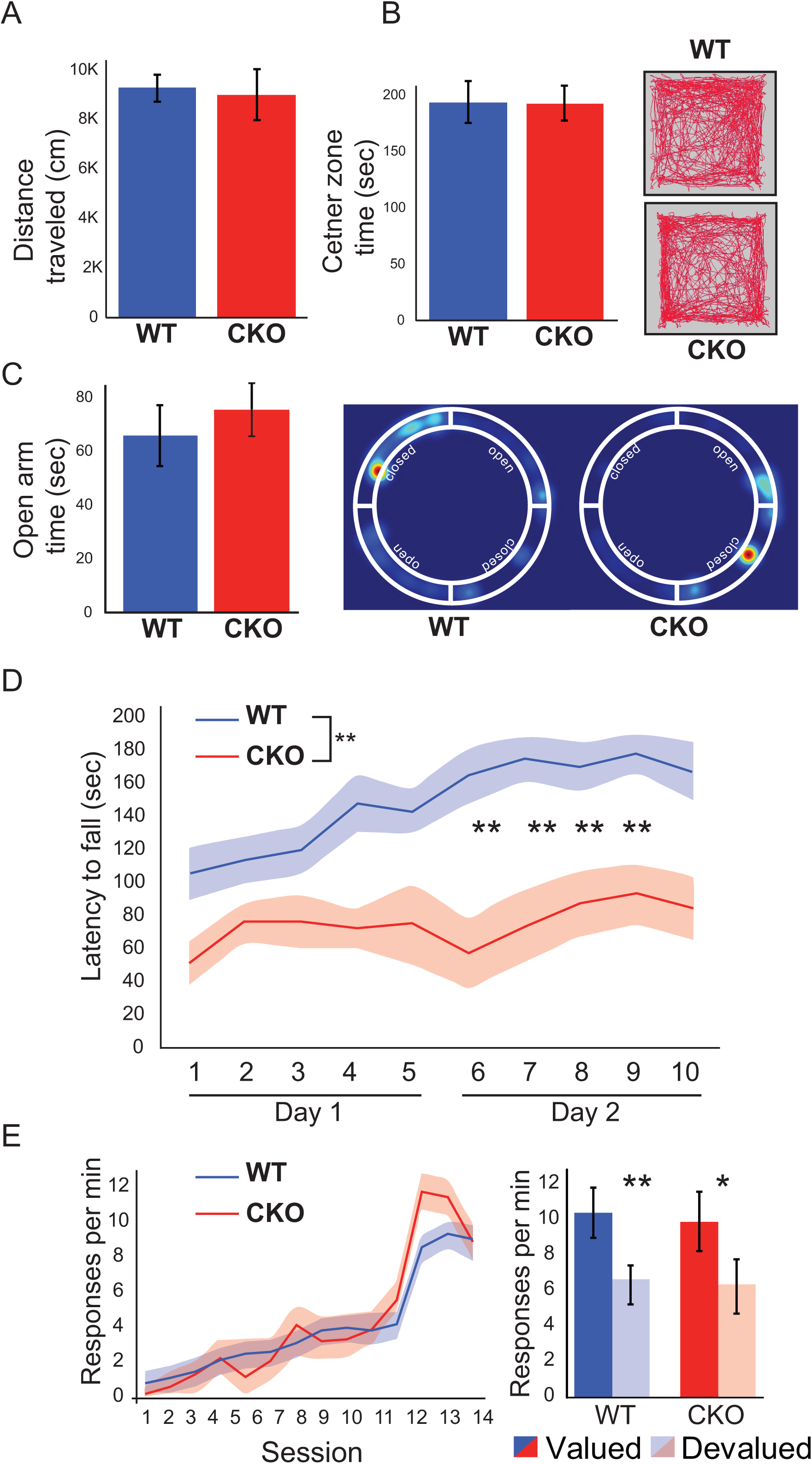
The effects of selective *Nrp2* deletion in adult layer V cortical neurons on sensorimotor and instrumental learning. **A-B**) No significant difference was detected between WT and *Nrp2* CKO mice for distance or time spent in the center of the open field test. **C**) No significant difference between WT and *Nrp2* CKO mice for time spent in open arm of the elevated zero maze. **D)** CKO mice showed a significant impairment in rotarod performance. Their overall latency to fall was signifianctly reduced, and *post-hoc* comparisons revealed significantly reduced latency on trials 6-9. **E)** CKO and control mice did not differ on acquistion of an instrumental response and performance across sessions with increasing response requirements. Following devaluation of the instrumental outcome, both groups of mice showed a significant reduction in response rate compared to when the outcome is still vaued. * *p* < 0.05; ** *p* < 0.005. Error bars represent ± 1 SEM.

We found that mice lacking Nrp2 in layer V cortical neurons were impaired on the accelerating rotarod, a sensorimotor learning task. We found an overall reduced latency to fall in *Nrp2* CKO mice compared to controls (ANOVA, F_1, 17_ = 16.25, *p* = 0.001) (Fig. 7D). Post-hoc comparisons with Bonferroni correction revealed significantly reduced latency to fall on trials 6-9 compared to controls (*p* < 0.005 for all comparisons). In contrast, we found no effect in these mice on instrumental goal-directed action. *Nrp2* CKO and littermate control mice acquired an instrumental response and showed similar patterns of responding across training sessions (Fig. 7E). During the extinction test following devaluation, both groups of mice significantly reduced their responses when the instrumental outcome was devalued compared to when the outcome was valued. This devaluation effect was significant in control mice (paired *t*-test, t_13_ = 3.46, *p =* 0.004) and *Nrp2 C*KO mice (*t*_13_ = 2.43 *p* = 0.03). Unlike the global *Nrp2* KO, selective deletion of Nrp2 in adult layer V cortical neurons did not affect goal-directed action control.

## Discussion

Our results demonstrate important roles for Nrp2 signaling in the development and maintenance of corticostriatal transmission and for striatal processing and function. We show that Nrp2 is present throughout the corticostriatal pathway (i.e. layer V cortical neurons, corticostriatal presynaptic axon terminals and striatal SPNs). We also demonstrated that the loss of Nrp2 disrupts corticostriatal transmission, increases SPNs excitability and spine density, and impairs goal-directed behavior. Furthermore, we found that acute deletion of Nrp2 selectively in layer V cortical neurons in adult animals increased spine density in apical dendrites of those neurons, altered corticostriatal plasticity, and impaired motor skill learning. Overall, our study uncovers novel and essential roles for Nrp2 signaling in adult corticostriatal synaptic transmission and associated behaviors.

### Nrp2 is expressed throughout the corticostriatal pathway and regulates dendritic spine frequency

Multiple areas of cerebral cortex, including sensory, motor, and association regions, project to the striatum (Goldman-Rakic and Selemon, 1986; Hintiryan et al., 2016; Jones et al., 1977; Kemp and Powell, 1970; Reiner et al., 2010; Royce, 1982; Shipp, 2017; Veening et al., 1980). This provides the striatum with the sensory and motor planning information needed for the basal ganglia to execute its role in motor control. The corticostriatal pathway mostly originates in the layer V (and to a lesser extent in layer 3) (Cowan and Wilson, 1994; Gerfen, 1989; Jones et al., 1977; Levesque et al., 1996; Parent and Parent, 2006; Reiner et al., 2010; Reiner et al., 2003; Wilson, 1987) and virtually all corticostriatal axons synapse on dendritic spines of SPNs.

As previously described Nrp2 is particularly enriched in layer V cortical neurons (Tran et al., 2009). In this study, we show for the first time that Nrp2 is also expressed in some presynaptic VGLUT1 positive corticostriatal axon terminals and in striatal SPNs at young postnatal ages and in adult mice. We previously showed that Sema3F and its receptor Nrp2 act as negative regulators of spine development and synapse formation (Tran et al., 2009). It has been shown that spine density and distribution are critical for proper functional synaptic transmission, integration and plasticity (Alvarez and Sabatini, 2007; Ballesteros-Yanez et al., 2006; Elston and DeFelipe, 2002; Nimchinsky et al., 2002). In this study, we found a significant increase in spine density in SPNs in adult *Nrp2^-/-^* mice consistent with the role of Nrp2 in negative regulation of spine formation during post-natal development.

### Nrp2 signaling mediates short-term facilitation of corticostriatal synapses and SPN excitability

We found that Nrp2 is expressed in corticostriatal axon terminals, which puts it in a position to modify presynaptic release mechanisms. Indeed, a similar role in presynaptic homeostatic plasticity has been described for Semaphorin 2b-Plexin B signaling (Orr et al., 2017). Consistently, loss of Nrp2 altered corticostriatal transmission, affecting neurotransmitter release probability and short-term facilitation which has been suggested to be important for the establishment and maintenance of membrane potential upstates in SPNs (Ding et al., 2008; Kasanetz et al., 2006; Wilson, 1993). Alternatively, the altered short-term facilitation we observed may be attributed to the increase in excitability in cortical neurons caused by loss of Nrp2 as it has been demonstrated that changes in excitability and presynaptic potentials at the level of the soma or even at the level of the dendrites can influence synaptic transmission (Christie and Jahr, 2008; Ludwar et al., 2009; Shu et al., 2006; Zbili et al., 2016). These two mechanisms are not mutually exclusive.

We also found that SPNs recorded *ex vivo* in adult *Nrp2^-/-^* mice exhibited a significant increase in excitability. These electrophysiological modifications in SPN’s in *Nrp2^-/-^*mice may influence spike integration, downstate to upstate transitions, and the induction of synaptic plasticity (Hernandez-Lopez et al., 2000; Nisenbaum et al., 1996; Shen et al., 2004; Wilson, 1995). These changes in SPNs’ excitability could be attributed to the loss of Sema3F-Nrp2 signaling within the striatum or they could be a consequence of upstream changes in layer V cortical activity and corticostriatal synapses. We addressed this question by acute deletion of Nrp2 specifically from cortical layer V neurons using *Nrpf/f;Etv1+/Cre* CKO mice treated with tamoxifen at adult age. In the *Nrp2* CKO mice, we found alterations to corticostriatal plasticity like those observed in *Nrp2^-/-^*mice, confirming that these changes in PPR are a presynaptic mechanism (Jackman and Regehr, 2017) involving the layer V cortical neurons. These results are also consistent with expression of Nrp2 in corticostriatal axons and with a role of Nrp2 in synaptic release of glutamate at corticostriatal synapses onto SPNs. Also, we demonstrated that in Nrp2 CKO mice, the intrinsic excitability of SPNs is not modified. These results suggest that expression of Nrp2 in SPNs is essential to maintain proper excitability of these cells.

### Nrp2 signaling influences motor learning and goal-directed action

We previously reported that *Nrp2^-/-^*mice are impaired in the accelerating rotarod, a motor learning task (Shiflett et al., 2015). Here we show that acute deletion of Nrp2 specifically in layer V cortical neurons is sufficient to impair rotarod behavior. Our result is consistent with previous findings demonstrating that motor skill learning critically depend on basal glutamate release and associated synaptic plasticity from corticostriatal terminals (Floyer-Lea and Matthews, 2005; Kupferschmidt et al., 2019; Yin et al., 2009b). We further demonstrate that *Nrp2^-/-^* mice are impaired on an instrumental task. *Nrp2^-/-^* mice were unable to adjust their responses to changes in the value of the instrumental outcome following specific satiety. *Nrp2^-/-^* mice showed reduced response rates during training, suggesting they had an impairment in learning. However, response rates during instrumental acquisition do not predict an animal’s choice strategy following devaluation. It is possible that *Nrp2^-/-^* mice and littermate controls learned similarly, and the deficit in *Nrp2^-/-^*following devaluation reflects an impairment in decision making.

Loss of goal-directed control occurs following lesions to the dorsomedial striatum and its afferent input from medial prefrontal cortex (Balleine et al., 2009; Ostlund and Balleine, 2005; Yin et al., 2005). Both of these structures are affected by loss of Nrp2. Synaptic plasticity within the dorsomedial striatum is necessary for goal-directed learning (Hawes et al., 2015; Shan et al., 2014; Shiflett et al., 2010) and likely underlies changes in activity patterns across the striatum as animals learn (Gremel and Costa, 2013; Hawes et al., 2015; Thorn et al., 2010; Yin et al., 2009a). Loss of Nrp2 may prevent goal-directed learning by interfering with corticostriatal synaptic plasticity, or by affecting SPN intrinsic excitability. Plasticity of intrinsic excitability in striatal neurons accompanies goal-directed learning (Daoudal and Debanne, 2003; Hawes et al., 2015). Therefore, the enhanced SPN intrinsic excitability we observed in *Nrp2^-/-^*mice may prevent intrinsic excitability changes necessary for learning.

While *Nrp2* CKO mice showed deficits in the motor skill learning, these mice showed intact performance in the instrumental goal-directed learning task contrasting with the results obtained in *Nrp2^-/-^* mice. One possible explanation for this difference is that motor learning and goal-directed learning differ in their rate of acquisition and may therefore rely on different forms of corticostriatal plasticity. During motor learning animals make trial-by-trial adjustments to motor patterns and these learning-related adjustments may depend on short-term corticostriatal synaptic plasticity of the type that is disrupted in the *Nrp2* CKO mice (Kupferschmidt et al., 2019). Goal-directed learning, in contrast, does not rely on trial-by-trial adjustment and may use a different set of synaptic-plasticity mechanisms that may be spared in CKO mice. Another possibility is that the acute deletion of Nrp2 in adulthood may not be sufficient to disrupt goal-directed learning, whereas Nrp2 deletion in layer V cortical neurons from a developmental stage may be necessary.

### Functional Implications

In sum, our findings have implications for understanding neurodevelopmental disease mechanisms. Alterations in the cellular anatomy and physiology of neurons comprising the corticostriatal pathway are found across many disorders, including Autism Spectrum Disorder, Schizophrenia, fragile X syndrome, and mental retardation (Hayrapetyan et al., 2014; Morris et al., 2015; Portmann et al., 2014; Schubert et al., 2015). Many of these diseases are accompanied by an increase in dendritic spine number and/or aberrant organization of synaptic connectivity (Barros et al., 2009; Carlson et al., 2011; Comery et al., 1997; Durand et al., 2012; Hayashi-Takagi et al., 2010; Hutsler and Zhang, 2010; Kim et al., 2013; Penzes et al., 2011; Ramakers, 2002). These disorders arise, in part, as a consequence of compromised excitation and inhibition (E/I) balance during brain development (Arnsten and Rubia, 2012; Brandon and Sawa, 2011; Geschwind and Levitt, 2007). Understanding how factors that contribute to the development and function of the corticostriatal pathway, such as we illustrate here with Nrp2, may ultimately lead to better treatments for these disorders.

## Acknowledgements

This study was supported by the NJ Governor’s Council for Medical Research and Treatment of Autism (CAUT17BSP022), NJDOH, the Rutgers – Newark Chancellor’s IMRT award to T.S.T, M.W.S. and J.M.T.,NIH NS034865 to J.M.T. and Rutgers University.

## Author Contributions

T.S.T and M.W.S conceived the study, M.A. performed the electrophysiological recordings, E.M. and C.E. performed the immunocytochemistry, C. E., A.K., D.E., and S.B. performed the behavioral tests, and K.V., F.S. assisted with some experiments and animal husbandry work. M.A., E.M., M.W.S., J.M.T. and T.S.T. contributed to the experimental design and interpretation of the results. M.A., M.W.S. J.M.T. and T.S.T. wrote the manuscript.

